# Evolutionary origins of archaeal and eukaryotic RNA-guided RNA modification in bacterial IS110 transposons

**DOI:** 10.1101/2024.06.21.599552

**Authors:** Hugo Vaysset, Chance Meers, Jean Cury, Aude Bernheim, Samuel H. Sternberg

## Abstract

Transposase genes are ubiquitous in all domains of life and provide a rich reservoir for the evolution of novel protein functions. Here we report deep evolutionary links between bacterial IS110 transposases, which catalyze RNA-guided DNA recombination using bridge RNAs, and archaeal/eukaryotic Nop5-family proteins, which promote RNA-guided RNA 2’-O-methylation using C/D-box snoRNAs. Based on conservation in the protein primary sequence, domain architecture, and three-dimensional structure, as well as common architectural features of the non-coding RNA components, we propose that programmable RNA modification emerged via exaptation of components derived from IS110-like transposons. Alongside recent studies highlighting the origins of CRISPR-Cas9 and Cas12 in IS605-family transposons, these findings underscore how recurrent domestication events of transposable elements gave rise to complex RNA-guided biological mechanisms.

## MAIN TEXT

Transposons are present in all domains of life and increase genetic diversity via horizontal gene transfer and genome rearrangements. Genes encoded by transposons have also been repeatedly repurposed and exapted during the evolution of essential cellular functions, ranging from V(D)J recombination^1^ and programmed genome elimination^2^, to RNA-centric processes such as pre-mRNA splicing^3^ and RNA-templated telomere maintenance^4^. In bacteria, RNA-guided DNA endonucleases encoded by CRISPR-Cas immune systems, including Cas9 and Cas12, were recently shown to have emerged from ancestral RNA-guided nucleases encoded by diverse transposons^5–7^. RNA-guided functions are also found in biological maintenance pathways such as RNA-guided RNA modification and splicing^8^, but their evolutionary origins are unknown.

Sequence-specific modification of ribosomal RNA (rRNA) is guided by abundant small non-coding RNAs in archaea and eukaryotes^9^ known as C/D-box small nucleolar RNAs (snoRNAs), which guide rRNA 2’-O-methylation to ensure accurate ribosomal processing and assembly^10^. The C/D-box snoRNAs contain evolutionarily conserved sequence elements, the C/D-box and the box C’/D’ motifs, which adopt kink-turns (K-turn). A similar K-turn structure is found in the U4 and U4atac small nuclear RNAs (snRNAs), which are required for spliceosomal assembly for sequence-specific pre-mRNA splicing and intron removal, a hallmark feature in nearly all eukaryotes^11–13^. In all cases, the K-turn is recognized by a L7Ae-family protein^14^ that then recruits secondary proteins containing a conserved Nop domain (Pfam PF01798) to the composite protein-RNA binding site^12^. Remarkably, we found that the origin of RNA-guided activities for both C/D-box snoRNAs and U4/U4atac snRNAs can be traced back to prokaryotic IS110 transposons, which similarly use guide RNA sequences, termed bridge RNAs, for programmable DNA recombination.

IS110 transposons are found in bacterial and archaeal genomes and encode a unique transposase/recombinase that features a DEDD active site, making it more similar to RuvC-like Holliday junction resolvases (HJR)^15^. While exploring IS110 Recombinase (IS110 Rec) homologs using HHpred^16^, we noticed an unexpected structural similarity to Nop5 and Prp31 proteins that both share a Nop domain, suggesting a potential shared ancestral relationship (**Supplementary Fig. 1**). The Nop domain is a key feature of archaeal Nop5 and its eukaryotic homologs, Nop56 and Nop58, which are structural subunits of C/D-box ribonucleoprotein (RNP) complexes that use snoRNA guides to direct precise 2’-O-methylation of target RNAs by the methyltransferase fibrillarin (**Fig. 1a**)^8^. Similarly, Prp31, an integral component of the U4/U6.U5 and U4atac/U6atac.U5 tri-snRNP complexes, contains a Nop domain to mediate its interactions with the RNP formed at the 5’ stem-loop of U4 or U4atac snRNP for spliceosomal assembly (**Fig. 1a**)^12,13,17,18^. Surprisingly, we noticed similar secondary structures when comparing bridge RNAs (IS110) and snoRNAs (C/D-box), including large bulged-out regions that function as paired guide RNA-like sequences (**Fig. 1a**). Based on this structural similarity, and on the recent discovery that IS110 Rec uses bridge RNA guides during DNA transposition^19,20^, we hypothesized that the archaeal and eukaryotic mechanisms for RNA-guided RNA modification may have originated from primordial RNA-guided DNA recombination processes found in IS110 elements, which later also became essential for RNA splicing.

**Figure 1.**
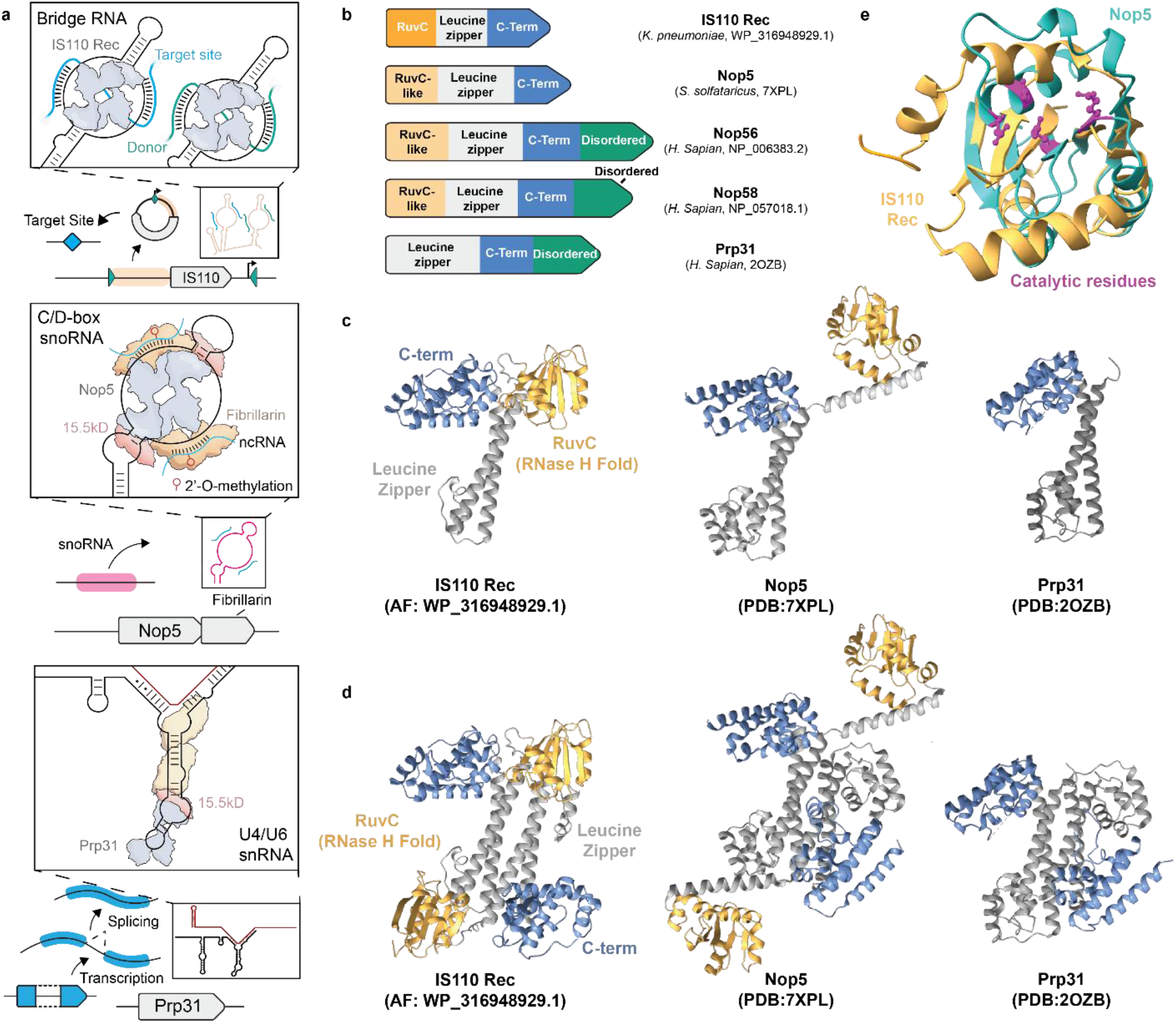
Functional and structural similarity between bacterial IS110 Rec, archaeal Nop5, and eukaryotic Prp31 proteins. **a,** Schematic illustrating RNA-guided mechanisms involving the IS110 Rec (top), C/D-box snoRNP (middle), and U4/U6 di-snRNP (bottom). IS110 bridge RNAs (black) direct site-specific transposition between donor (green) and target (blue) DNA molecules via RNA-DNA base-pairing interactions and catalytic activity of the IS110 Rec. C/D-box snoRNPs use similar structured RNAs to direct 2’-O-methylation via RNA-RNA base-pairing interactions, with Nop5 playing a key structural role in the assembly of the RNP. A ribonucleoprotein complex formed at the 5′-stem loop of U4/U6 di-snRNAs exploits Prp31 to stabilize the U4/U6 duplex, required for the assembly of the spliceosome. **b,** Comparison of IS110 Rec, Nop5, Nop56, Nop58, and Prp31 protein domain architectures. Both IS110 Rec and Nop5 proteins exhibit a similar arrangement of domains, including an N-terminal RuvC-like domain followed by a leucine zipper domain and C-terminal region rich in α-helices **c,** Structural comparison of Nop5 (PDB: 7XPL) and Prp31 monomers (PDB: 2OZB) with an AlphaFold (AF) prediction of IS110 Rec. Domains are colored according to the domain annotations in **b**. The C-terminal disordered region of Nop56/Nop58 and Prp31 is not displayed, for clarity. **d,** Structural comparison of dimeric forms of IS110 Rec, Nop5 (PDB: 7XPL), and Prp31 (PDB: 2OZB), colored as in **c**; the IS110 Rec structure is an AlphaFold multimer prediction. **e,** Sequence-guided structural alignment of the AlphaFold-predicted IS110 Rec (orange; WP_316948929.1) with Nop5 (shown in teal; PDB: 7XPL), focusing on the N-terminal RuvC-like region. The active site region is highlighted in purple, with catalytic residues of IS110 Rec shown. The RMSD between 23 pruned atom pairs is 1.093 Å.

We investigated these similarities further using AlphaFold2^21^, to compare the predicted tertiary structure of IS110 Rec with experimental structures of Nop5 and Prp31. These analyses revealed pronounced structural homology, with IS110 Rec and Nop5 sharing the same overall architecture, including an N-terminal RuvC-like domain followed by a leucine zipper/coiled-coil region and C-terminal α-helical domain; Prp31 exhibits a similar fold but lacks the N-terminal domain (**Fig. 1b-d**). The presence of a catalytically inactivated RNaseH/RuvC-like fold in Nop5 has not been reported before, and this observation, together with the insertion of the alpha-helix that repositions the domain relative to the leucine zipper domain, highlights a potential important repurposing event that selected for fibrillarin recruitment rather than nucleic acid cleavage (**Fig. 1e**, **Supplementary Fig. 2a**). All three proteins feature α-helical bundles within the C-terminal domain, though Nop5 and Prp31 both lack the β-sheet that forms the catalytic serine active site essential for IS110 transposition (**Supplementary Fig. 2b**)^19,20^. Interestingly, AlphaFold Multimer models predict IS110 oligomerization via its coiled-coil region, similar to Nop5, which forms dimers that act as a scaffold for the snoRNP complex^22–24^ (**Fig. 1d**). This multimeric propensity hints at more complex structural arrangements (**Supplementary Fig. 3**), akin to the higher-order organization commonly observed in other serine recombinases families^25^.

To systematically identify all the homologs of the Nop family of protein, we used sensitive searches with either Foldseek^26^ or HHpred^27^. We identified a set of homologous proteins that includes archaeal Nop5 and eukaryotic Nop56/Nop58 homologs, eukaryotic Prp31, and bacterial IS110 Rec (**Methods, Supplementary Tables 1-2**). More systematic homology searches against a database of 30,255 prokaryotic and eukaryotic genomes failed to reveal homologs outside these three families (**Supplementary Table 3**), suggesting an absence of additional proteins that exhibit this particular domain composition. When we analyzed the N-terminal domain of the IS110 Rec in isolation, homologs within the family of RuvC-like HJR were identified (**Supplementary Table 3**)^28,29^. Homology searches using HHpred starting from the Nop5- and Prp31-encoded Nop domain alone robustly retrieved the C-terminal region of the IS110 Rec (Pfam PF02371, probability=0.96, **Supplementary Table 4**), and a reciprocal search starting from the ‘Nop’ domain of the IS110 Rec similarly returned the homologous domains from Nop5 and Prp31, with high confidence (Pfam PF01798, probability=0.96, **Supplementary Table 4**). Collectively, these complementary lines of evidence provide strong support for a shared ancestry linking the IS110 Rec with Nop5- and Prp31-family proteins.

We then investigated whether the Nop5 family emerged from IS110 Rec. Testing this hypothesis required constructing a phylogeny to evaluate the evolutionary relationship between IS110 Rec and Nop5, relative to other protein families. As sensitive full-length protein homology searches did not yield homologs beyond the IS110 Rec, Nop5 and Prp31 families, we focused on each individual protein domain (N-terminal, leucine zipper, C-terminal), retrieved all corresponding homologs, built domain-level alignments, and then concatenated them to construct a phylogeny (**Fig. 2a**). This method allowed us to capture the evolutionary history of bacterial/archaeal IS110 Rec, archaeal/eukaryotic Nop5, and all their domain-level homologs within a single phylogenetic tree (**Methods**). The obtained alignment revealed broad conservation of key residues across the Nop domain, including three highly conserved glycine residues in the C-terminal α-helical domain (**Supplementary Fig. 4a,b**) and an arginine residue that interact with the backbone of RNA^22^. Both IS110 Rec and Nop5 form monophyletic clades, and the position of Nop5 and Rec clades within the RuvC-like superfamily indicates that the two protein families are close relatives, rather than Nop5 following a unique evolutionary trajectory to converge on the same domain architecture (**Supplementary Fig. 5**). Interestingly, the structure of the phylogenetic tree indicates that archaeal and eukaryotic Nop5 homologs did not descend from an archaeal IS110 Rec clade present in our database, and instead suggests that the Nop5 lineage either originated from an archaeal IS110 Rec clade that is extinct today, or from a more ancient IS110 Rec ancestor (**Fig. 2c**, **Supplementary Fig. 5**). This scenario is also supported by full-length protein and single-domain trees including Rec and Nop5 homologs (**Supplementary Fig. 6–7**). Given the widespread distribution of IS110 transposons in bacteria and the diverse changes in Nop5 (notably loss of the RuvC active site), the IS110 Rec is likely to represent the ancestral state. Finally, we exploited the shared Nop domain to build a phylogenetic tree of archaeal Nop5, eukaryotic Nop56/Nop58, and Prp31, which revealed the likely emergence of Prp31 from a Nop5 duplication event that likely occurred in early eukaryotes (**Fig. 2b,c**)^30^.

**Figure 2.**
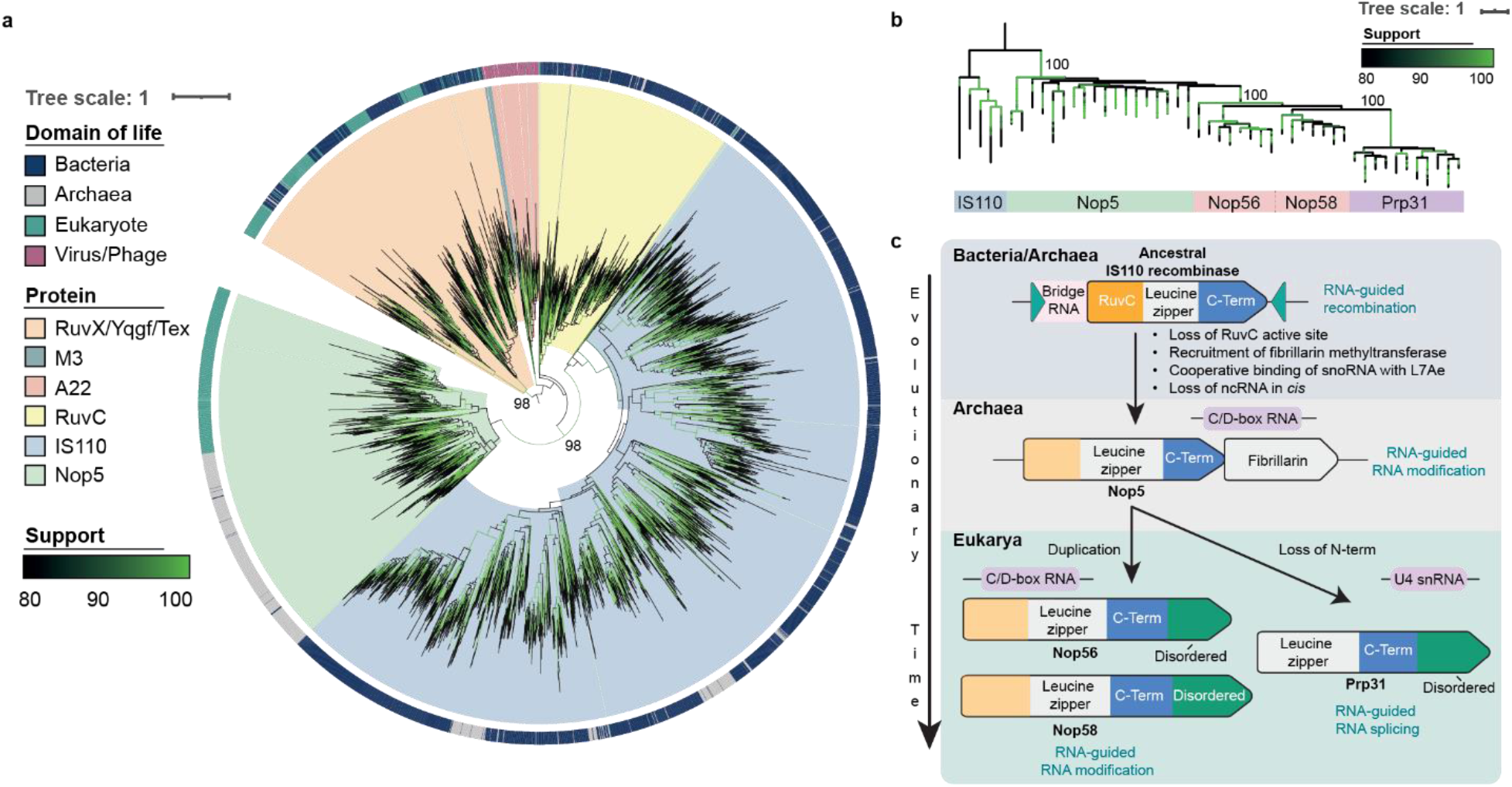
IS110-family recombinases share a common evolutionary ancestor with Nop5-family proteins involved in snoRNP assembly. **a,** Phylogenetic tree including IS110 Rec, Nop5, Nop56/Nop58, and RuvC-like homologs (RuvC, A22, M3, RuvX/YqgF/Tex) that were included to help with assessing evolutionary relationship. The three protein domains conserved in IS110 Rec and Nop5 were separately aligned, and multiple sequence alignments were then concatenated (see **Methods**). The domain of life origin for each homolog is denoted in the outer ring, and UltraFast Bootstrap (UFBoot) values are shown in shades of green. **b,** Phylogenetic tree built on the Nop domain of Nop5 (archaea, green), Nop56 and Nop58 (eukaryotic, pink), and Prp31 (eukaryotic, purple) (see **Methods**). The tree is rooted on representative sequences of the IS110 Rec (blue) and subsampled to enhance readability; the full version is presented in **Supplementary Fig. 5c**. UFBoot support values for each branch are indicated in green and are explicitly specified for key branches. **c,** Parsimonious evolutionary timeline explaining the emergence of Nop5 and Prp31 across the tree of life, from their last common ancestor with the IS110 Rec. The Nop5 lineage likely originated in early archaea from an ancestral protein that resembled the IS110 Recombinase currently found in both bacteria and archaea. Nop5 proteins diverged from the IS110 Rec through multiple events of gain/loss of specific biochemical properties, and the transition of bridge RNAs to C/D-box and U4 snRNAs. The Prp31 lineage likely emerged in early eukaryotes via a Nop5 duplication event and further divergence through loss of the N-terminal domain.

Collectively, our findings provide strong support for the emergence of both Nop5 and Prp31-family proteins from ancestral IS110 Rec (**Fig. 2c**), unifying the seemingly disparate processes of RNA-guided DNA recombination, RNA-guided RNA modification, and RNA-guided RNA splicing. We propose that the Nop5 lineage likely emerged in early archaea from an ancestral protein which probably possessed similar structural and functional features as the IS110 Rec present in extant bacteria and archaea. Nop5 evolution was marked by significant functional changes, including loss of nuclease activity in the N-terminal domain and the acquisition of protein-interacting functions to engage other components of snoRNP/snRNP complexes for novel RNA-guided processes. When taken together with recent work demonstrating a transposon origin for RuvC nuclease domain-containing CRISPR effectors such as Cas9 and Cas12^5,6,31,32^, our work highlights the repeated emergence of complex RNA-guided functions in bacterial mobile genetic elements, showcasing their pervasive influence on complex cellular pathways. These diverse, programmable features have been exapted by their host organisms to fulfill novel biological roles and present exciting opportunities to innovate and advance genome engineering technologies.

## METHODS

### Database of eukaryotic, archaeal and bacterial genomes

4,616 eukaryotic genomes from 2,407 species and 22,804 bacterial complete genomes were downloaded from Genbank in January 2022^33^. 2,339 archaeal genomes were downloaded from the Genome Taxonomy Database (GTDB) in January 2023 and concatenated with 496 in-house Asgard archaea genomes, resulting in a database of 2,835 archaeal genomes^34^. Altogether, our database comprised 30,255 genomes across the three domains of life.

### Remote homology between IS110 Rec and Nop5

Remote homology detection was assessed using the HHpred webserver and Foldseek (version 8-ef4e960)^26,27^. The initial observation suggesting a putative homology link between Nop5 and the IS110 Rec was made by querying individual IS110 Rec sequences from *K. pneumoniae* (A0A809T667) and *S. flexneri* (QKW72470) against the Pfam-A (version 36) and the PDB_mmCIF70 databases using the HHpred, with default parameters (**Supplementary Table 2**).^27,35^. This association was also assessed using Foldseek, querying (options -c 0.8 and --max-seqs 10000) representative sequences of IS110 Rec, Nop5, Nop56, and Nop58 against the UniProt50 Foldseek database, as downloaded in February 2024^26,36^. The obtained hits were filtered (probability > 0.8) and further annotated using the Pfam-A (v36) database to determine their identity (**Supplementary Table 1**).

### Protein structure prediction

When experimentally solved protein structures were not available from the RCSB Protein Data Bank, protein structure predictions were retrieved from the AlphaFold Protein Structure database (AFDB)^36,37^. When neither experimental structures nor predicted protein structures were readily available in these databases, protein structure prediction was performed using AlphaFold2 through the ColabFold distribution (version 1.5.2) as implemented in the localcolabfold package^21,38^. When predicting multimeric structures, AlphaFold-Multimer v3 was used^39^. Protein structure prediction was performed using three independent models with 20 prediction recycles each. Template information was used, and multiple sequence alignments were computed in mmseqs2_uniref_env mode using only paired sequences. Protein structures were visualized and drawn using ChimeraX (version 1.7.1)^40^.

### Systematic identification of IS110 Rec and Nop5 homologs

IS110 Rec homologs were identified using iterative profile HMM search on our database of 30,255 bacterial, archaeal and eukaryotic genomes. An initial BLAST (version 2.12.0) search was performed using a *K. pneumonia* IS110 Rec (A0A809T667) as a query, in order to identify all closely related homologs (-max_target_seqs 1000000 -evalue 1e-80)^41^. The retained hits (n=6,407) were aligned using the MAFFT (version 7.520) L-INS-i algorithm (--localpair), and an initial profile HMM was built and used as a query to search for additional homologs using hmmsearch from the HMMER package (version 3.3.2), with default parameters^42,43^. All hits (n=55,007) with an e-value below 1e-5 and a protein length between 200–1,000 amino acids were retained for another round of search. Hits were clustered at 80% identity and 80% coverage using mmseqs easy-cluster (version 13.45111)^44^. Representative sequences (n=4,047) were aligned using MAFFT L-INS-I, and a new profile HMM was built from this alignment. All multiple sequence alignments (MSA) were visualized using the Unipro UGene software (version 49.1)^45^.

The same procedure was repeated to obtain 58,533 hits satisfying the inclusion criteria (e-value < 1e-5 and 200 < length < 1,000). Homology search was then terminated, as another round of search did not yield any new hits satisfying the inclusion criteria. The set of identified IS110 Rec homologs was further filtered to include only proteins hits covering at least 80% of the profile HMM, leading to a final set of 58,533 IS110 Rec homologs among bacteria and archaea (**Supplementary Table 3**). As proteins retrieved using this procedure were mostly of bacterial origin, an extra round of search was performed using a profile HMM built from only the identified archaeal IS110 Rec as a query (n=662), against the database of archaeal genomes. This round of search did not yield any additional archaeal IS110 Rec homologs.

Nop56 and Nop58 homologs were identified using a similar iterative profile HMM search procedure. Animal orthologs of Nop56 and Nop58 were retrieved from the NCBI Orthologs database and clustered at 98% identity and 98% coverage using mmseqs, leading to a set of 498 representative orthologs which were aligned using MAFFT L-INS-i to build an initial Nop56/Nop58 profile HMM^42,44,46^. Homology search using this profile led to 15,253 hits satisfying the inclusion criteria (e-value < 1e-5 and 300 < length < 700 amino acids), which included the archaeal Nop5 protein and the eukaryotic proteins Nop56, Nop58, and Prp31. Those homologs were clustered at 80% identity and 80% coverage, and representative sequences (n=2,397) were used for another round of search against the genome database. This round yielded 15,932 hits satisfying the inclusion criteria (e-value < 1e-5 and 300 < length < 700 amino acids) (**Supplementary Table 3**). Hits were reannotated using Pfam-A (v36), and all belonged to either the Nop5/Nop56/Nop58 or Prp31 family.

### Systematic identification of individual domain homologs

We sought to identify candidate homologs of each individual domain for IS110 Rec and Nop5. We decided to split the proteins into three parts based on structural features: (i) the N-terminal domain (RuvC-like in the case of IS110 Rec), (ii) the leucine zipper domain, and (iii) the C-terminal domain. Each domain was manually extracted from the whole protein alignments in order to build domain-level profile HMM and search for putative homologs in our database. First, single protein profile HMM were built and used as queries in hmmsearch against our database of genomes^43^. As no significant hits belonged to new protein families using this approach (e-value < 1e-5), ‘mixed’ profile HMM were built by aligning sequences of a domain shared between distinct proteins (e.g. both IS110 Rec and Nop5 N-terminal domains). Again, no novel significant hits could be identified using this approach.

In order to increase the sensitivity of the homology search for each individual domain, a MSA including the N-terminal domain of representative sequences (mmseqs easy-cluster at 60% identity and 60% coverage) of both IS110 Rec and Nop5, as well as of the compact RuvC nuclease domain, was queried in HHpred against the Protein Data Bank and the Pfam-A databases (5 iterations against UniRef30 for MSA generation, minimum probability of 20%)^47^. Hits that satisfied inclusion criteria were filtered (probability > 0.8) for further analysis. This approach uncovered new proteins from the family of the RuvC-like Holliday junction resolvase, in particular RuvC (PF02075), poxvirus A22 (PF04848), *Lactococcus* phage protein M3 (PF07066), and the YqgF/RuvX/Tex family of proteins (CL0580 / PF14639, PF03652, PF16921). Proteins from these families were retrieved in our database by querying their respective Pfam-A profile HMM in hmmsearch (with option --cut_ga) against our database of genomes^35,43^. Those hits were added to the list of candidate homologs for the N-terminal domain of Nop5 and IS110 Rec, for downstream phylogenetic inference. The same approach was applied to the other domains of Nop5 and IS110 Rec but did not provide any significant matches apart from Nop5 and IS110 Rec themselves. Foldseek searches were also performed using the individual domains of representative sequences for IS110 Rec, Nop5, Nop56, Nop58, and Prp31. Individual domains were manually extracted for these proteins and queried by Foldseek against the UniProt50 database (options -c 0.8 and – max_seqs 10000)^26,36^. Hits were filtered (probability > 0.8) and annotated using profile HMM from the Pfam-A database^35,43^ (**Supplementary Table 1**).

### Phylogenetic inference regarding IS110 Rec, Nop5, RuvC, and RuvC-like homologs

We first built a phylogenetic tree of full-length IS110 Rec and Nop5 proteins (**Supplementary Fig. 4**). This tree was used to infer specific subfamilies within the IS110 Rec and Nop5 families and to assess the relationship between the IS110 Rec and Nop5 proteins. All IS110 Rec and Nop5 proteins detected in the database of genomes were first clustered at 60% identity and 60% coverage using mmseqs easy-cluster^42^. Representative sequences (n=2,898) were aligned using MAFFT L-INS-I, and the positions with more than 99% gaps were trimmed from the MSA using Clipkit (version 1.3.0, -m gappy -g 0.99), leading to a trimmed alignment of 1,237 positions^42,48^. IQTree was used to build a maximum likelihood phylogenetic tree that was optimized for 900 iterations using the Q.pfam+G4 substitution model, and branch support was assessed using 1000 UltraFast Bootstrap (UFBoot) trees^49–51^.

Individual protein domains (N-terminal, leucine zipper, and C-terminal) were separately aligned using MAFFT L-INS-i, including representative sequences (mmseqs easy-cluster at 60% identity and 60% coverage) of the proteins that possess each domain^42,44^. These alignments were first used to build domain-level phylogenetic trees for the N-terminal and C-terminal domains, including all relevant homologs for each domain (**Supplementary Fig. 6**). Single domain phylogenies were built in order to assess putative evolutionary convergence involving multiple independent domain assembly events which would translate in incongruent domain-level trees. For the N-terminal domain, we included RuvC and RuvC homologs (RuvC, A22, M3 and YqgF/RuvX/Tex), in addition to IS110 Rec and the Nop5 N-terminal domain^28,29^. Proteins were clustered at 60% identity and 60% coverage using mmseqs, and representative sequences were aligned using MAFFT L-INS-i^42,44^. The MSA was manually inspected and trimmed using Clipkit^32^. Maximum likelihood trees were searched for 3000 iterations using IQTree (version 2.2.0.3), with the Q.Pfam+G4 substitution model^33,35^. Branch support was assessed by generating 1000 UFBoot trees^50^. For the C-terminal domain, only IS110 Rec and Nop5 proteins were included, as no other candidate homologs were identified. Sequences were processed as described for the N-terminal domain (clustering at 60% identify and 60% coverage using mmseqs easy-cluster, alignment using MAFFT L-INS-I, and trimming using Clipkit gappy to positions containing more than 99% gaps), and a maximum likelihood tree was inferred using IQTree for 3000 iterations with the LG+G4 substitution model^42,44,48,49,51^. Branch support was assessed using 1000 UFBoot trees^50^. The MSA was also visualized using UGene, and conserved residues were mapped onto predicted or experimental structures of IS110 Rec and Nop5, respectively, using ChimeraX^40^.

The domain-level alignments were also used to build a phylogenetic tree at the full-length protein level for IS110 Rec and Nop5 (**Fig. 2a**). To do so, individual domain alignments were concatenated to obtain a unique MSA modeling the full-length protein, including representative sequences (clustered at 60% identity and 60% coverage using mmseqs) of each domain-level homolog identified during the search (n=3,950 sequences in total), including members of the RuvC-like family for the N-terminal domain of IS110 and Nop5^44^. This strategy was adopted as (i) aligning at the single domain-level allows to obtain good quality MSA including the homologs possessing each individual domain in the absence of full protein homologs and (ii) building a tree of the concatenated MSA allows to infer the evolutionary history of the full IS110 Rec and Nop5 proteins in a single tree. The MSA was trimmed using clipkit gappy (removing positions with more than 99% gaps), leaving 762 positions satisfying the inclusion criteria^48^. A maximum likelihood phylogenetic tree of the full-length protein was then inferred from this concatenated MSA using IQTree for 4000 iterations with the Q.Pfam+G4 substitution model and 1000 UFBoot trees for assessing branch support^44,49–51^. Among all the included families of proteins (RuvC, YqgF/RuvX/Tex, A22, M3, IS110 Rec, and Nop5), the YqgF/RuvX/Tex was chosen as an outgroup to root the tree because of its more important divergence to other RuvC-like proteins, as can be seen in its catalytic residues (DED instead of DEDD in all other RuvC-like proteins, including IS110 Rec)^28,29^.

To assess the phylogenetic relationship between the Nop5 and Prp31 lineages, we built a phylogenetic tree of the Nop domain from Nop5 and Prp31 (**Supplementary Fig. 5**). The Nop domain refers to the section of the protein spanning from the α-helical bundle separating the two α-helices of the leucine zipper region, to the C-terminal orthogonal α-helical bundle, including the second helix of the leucine zipper. This region was chosen to build the phylogenetic tree of the proteins, as it contains conserved residues among Nop5/Nop56/Nop58 and Prp31; in contrast, the N-terminal RuvC-like domain of Nop5 proteins is not present in Prp31. The Nop domain region was manually extracted from all identified Nop5 and Prp31 homologs, and proteins were clustered at 80% identity and 80% coverage using mmseqs easy-cluster^44^. Representative sequences (n=1,703) were aligned using MAFFT L-INS-I, and a maximum likelihood tree was inferred using IQTree (LG+G4 substitution model, 5000 iterations, 1000 UFBoot trees)^44,49–51^.

## Data availability

All code and raw data are available on the GitHub repository of the project (https://github.com/mdmparis/is110_nop5)

## ACKNOWLEDGEMENTS

We thank Eduardo Rocha, Homa Ghalei, and members of the Sternberg lab for carefully reading the manuscript, providing constructive feedback, and engaging in fruitful discussions. Several bioinformatic analyses were performed on the Core Cluster of the Institut Français de Bioinformatique (IFB) (ANR-11-INBS-0013). C.M. was supported by an NIH F32 Postdoctoral Fellowship (GM143924) and an ASGCT Career Development Award. A.B., J.C. and H.V. were supported by an ERC Starting Grant (PECAN 101040529). S.H.S. was supported by NSF Faculty Early Career Development Program (CAREER) Award 2239685, a Pew Biomedical Scholarship, an Irma T. Hirschl Career Scientist Award, and a generous startup package from the Columbia University Irving Medical Center Dean’s Office and the Vagelos Precision Medicine Fund.

## AUTHOR CONTRIBUTIONS

C.M. and S.H.S. conceived of the project. C.M. and H.V. performed protein structural predictions and analyses. A.B., J.C. and H.V. designed and interpreted systematic homology searches and phylogenetic analyses, which were performed by H.V. All authors discussed the data and wrote the manuscript.

## SUPPLEMENTARY FIGURES

**Supplementary Figure 1.**
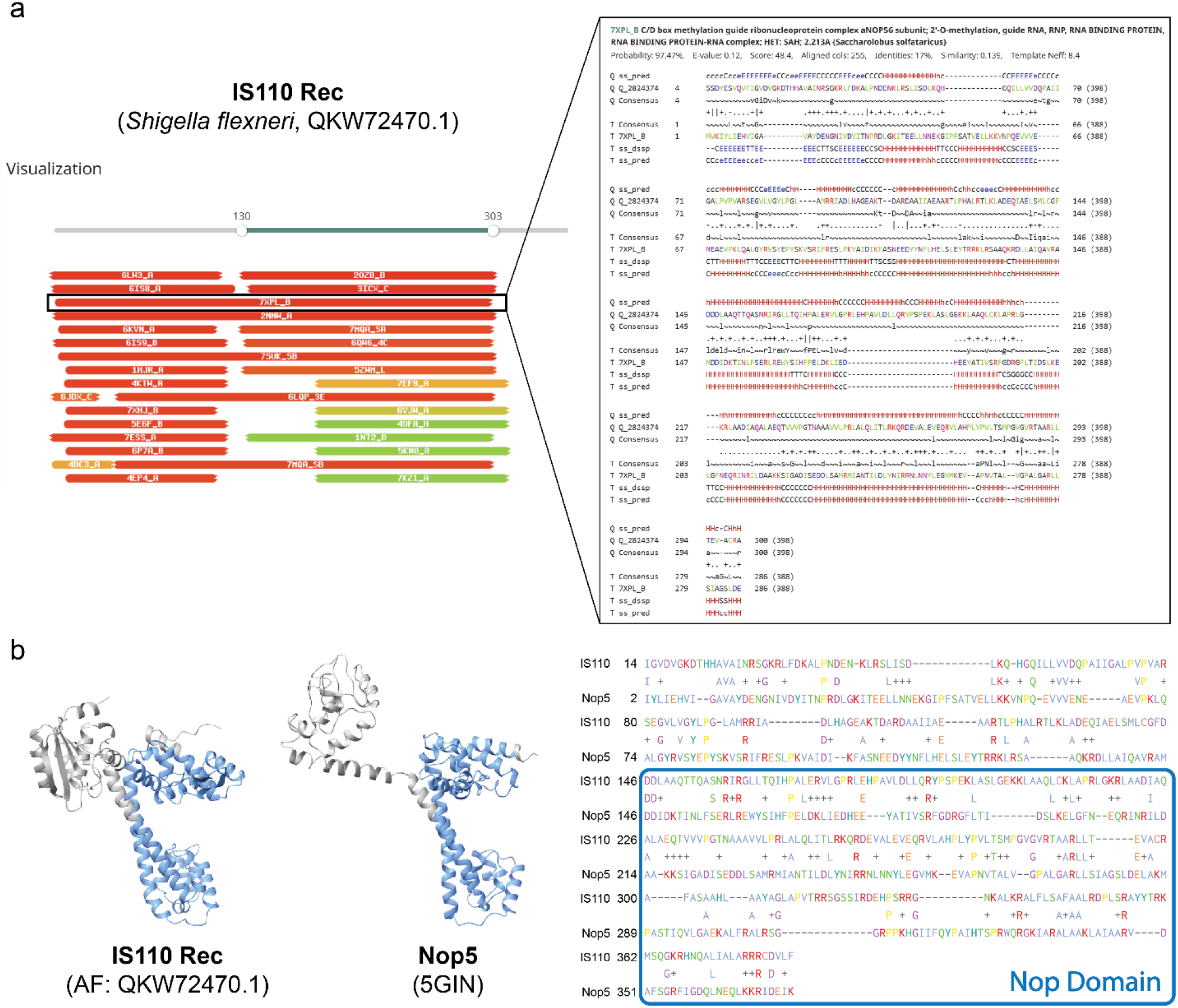
Profile HMM-based homology detection and secondary structure prediction of IS110 transposase. **a,** Homology detection and secondary structure prediction for IS110 Rec were performed using HHpred. The left panel displays the alignment of IS110 Rec with various protein domains identified through HHpred, indicating structural and functional similarities based on structural hits from the Protein Data Bank (PDB). The right panel provides a detailed view of the alignment with C/D-box snoRNP subunit Nop5 from *Sulfolobus solfataricus* (7XPL_B), showing the probability of homology, E-value, and secondary structure elements. Predicted secondary structures (α-helices and β-strands) are indicated above the amino acid sequence. **b,** Homology detection and secondary structure prediction for IS110 Rec were performed using Foldseek. The left panel displays the AlphaFold-predicted structure of IS110 Rec (QKW72470.1) and the empirically determined structure of Nop5 from *Sulfolobus solfataricus* (5GIN). The Nop domain is outlined in blue. 3Di+AA FoldSeek alignment are shown on the right, with the Nop domain outlined in blue.

**Supplementary Figure 2.**
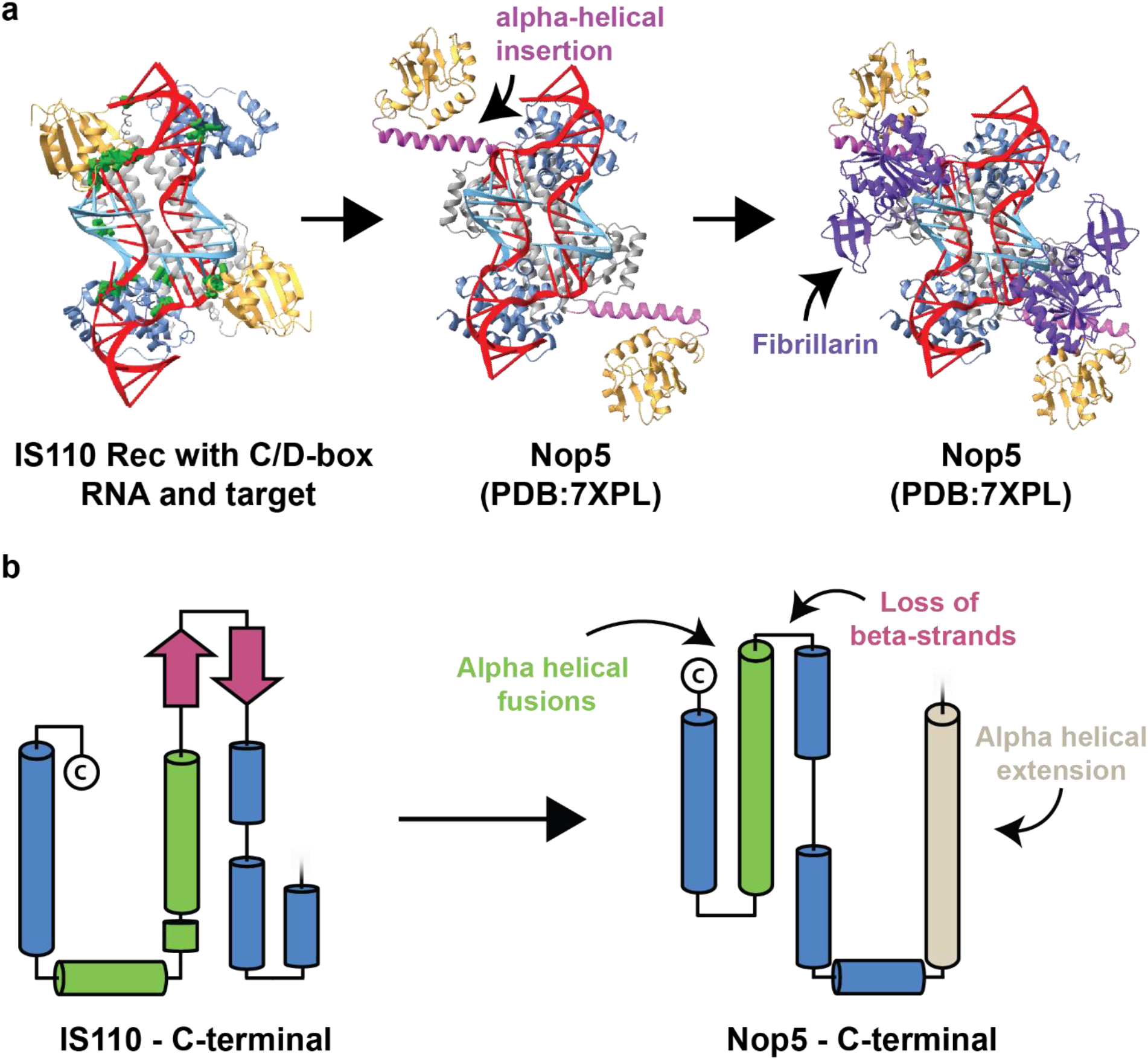
Key structural differences between the IS110 Rec and Nop5. **a,** An AlphaFold2 structure prediction of a bacterial IS110 Rec (WP_316948929.1) dimer overlaid with an archaeal C/D-box RNA (left) exhibits limited steric clashing, indicated in green; the RuvC-like nuclease domains (orange) are positioned in close proximity to the target nucleic acid sequence (light blue). The middle panel displays an experimental structure of Nop5 bound to C/D-box RNA, highlighting an α-helical insertion (pink) that displaces the N-terminal RuvC-like fold, as compared with IS110 Rec. The right panel illustrates that the fibrillarin methyltransferase, shown in purple, is positioned within this region to gain access to the target nucleic acid for 2’-O-methylation. **b,** Graphical representation of the IS110 Rec C-terminal domain secondary structure, as predicted by AlphaFold (left), with α-helices represented by barrels and β-strands by arrows. On the right, the graphical representation of the Nop5 C-terminal domain is shown (PDB: 7XPL), with changes relative to IS110 Rec highlighted.

**Supplementary Figure 3.**
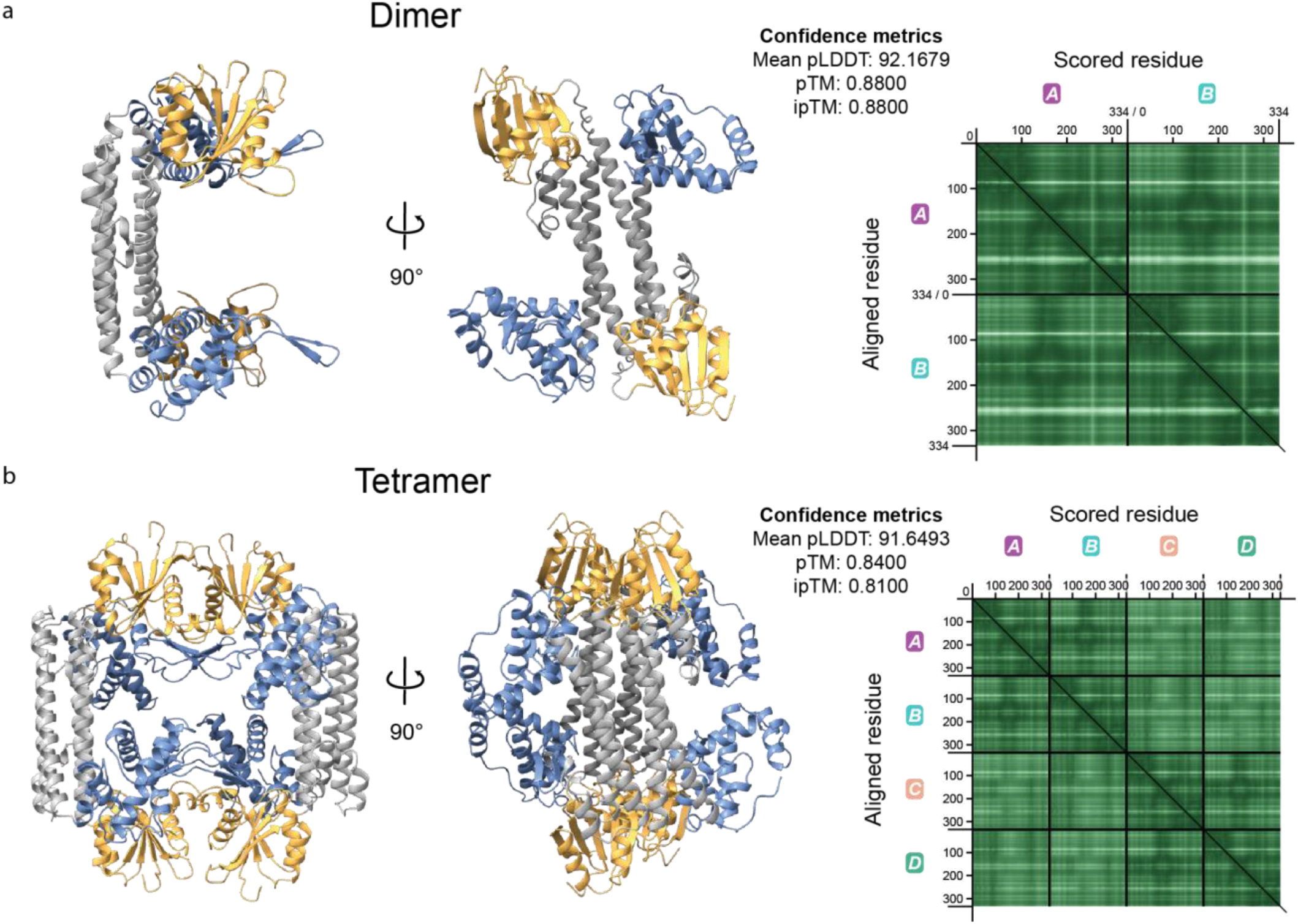
Predicted oligomeric structures of IS110 Rec by AlphaFold2. **a,** AlphaFold2 multimer structural predictions of an IS110 Rec dimer (WP_316948929.1, left). The proteins are colored by domain, as in Fig. 1b, with RuvC domain in orange, leucine zipper in gray, and C-terminal domain in blue. Confidence metrics are accompanied by PAE matrices for each monomer (right), indicating a high-confidence prediction of dimer formation. Predicted Aligned Error (PAE) matrices quantify the expected positional error for each residue in a predicted protein structure relative to its counterpart in the true structure when aligned. **B,** AlphaFold-multimer structural predictions of an IS110 Rec tetramer, shown as in **a.**

**Supplementary Figure 4.**
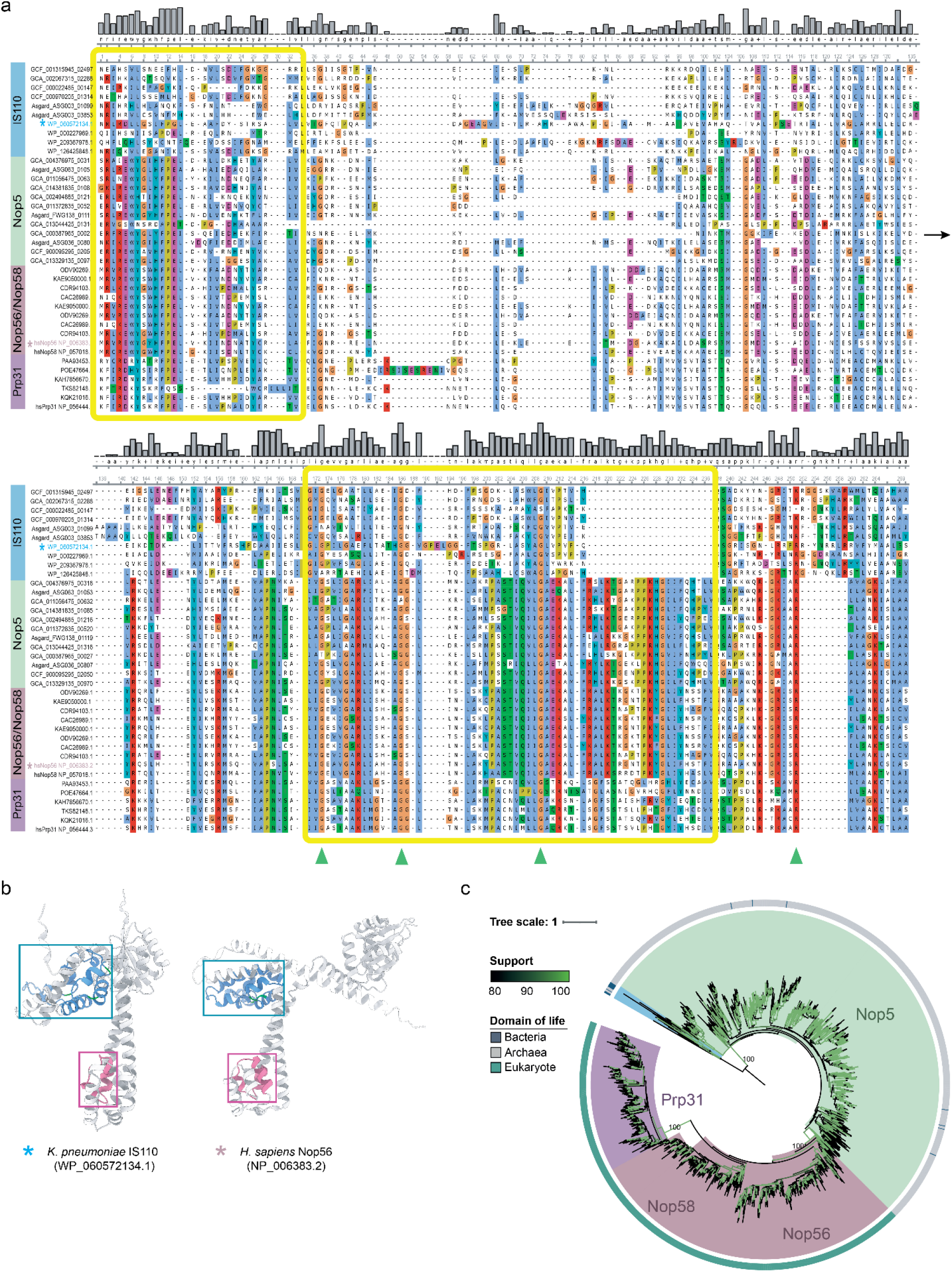
Alignment and phylogenetic tree built on the Nop domain of IS110 Rec, Nop5, Nop56/Nop58, and Prp31 homologs. **a,** Multiple sequence alignment (MSA) of the Nop domain from representative IS110 Rec (blue), Nop5 (green), Nop56/Nop58 (pink), and Prp31 (purple) homologs. Two regions exhibiting moderate to strong amino acid conservation are highlighted by colored boxes (yellow), and green triangles indicate conserved residues. Colored stars (light blue and pink) near protein names indicate proteins whose structures are highlighted in panel **b. b,** AlphaFold2 model of representative IS110 Rec (WP_060572134.1) and Nop56 proteins (NP_006383.2). Regions highlighted in pink and blue correspond to the boxed regions in **a**, and residues highlighted in green correspond to three highly conserved glycines and an arginine. **C,** Phylogenetic tree built on the Nop domain of Nop5, Nop56/Nop58, and Prp31, with a set of representative IS110 Rec homologs chosen as an outgroup to root the tree (blue). UFBoot support values for each branch are indicated in green and are explicitly specified for key branches. A subsampled version of this tree is presented in Fig. 2b.

**Supplementary Figure 5.**
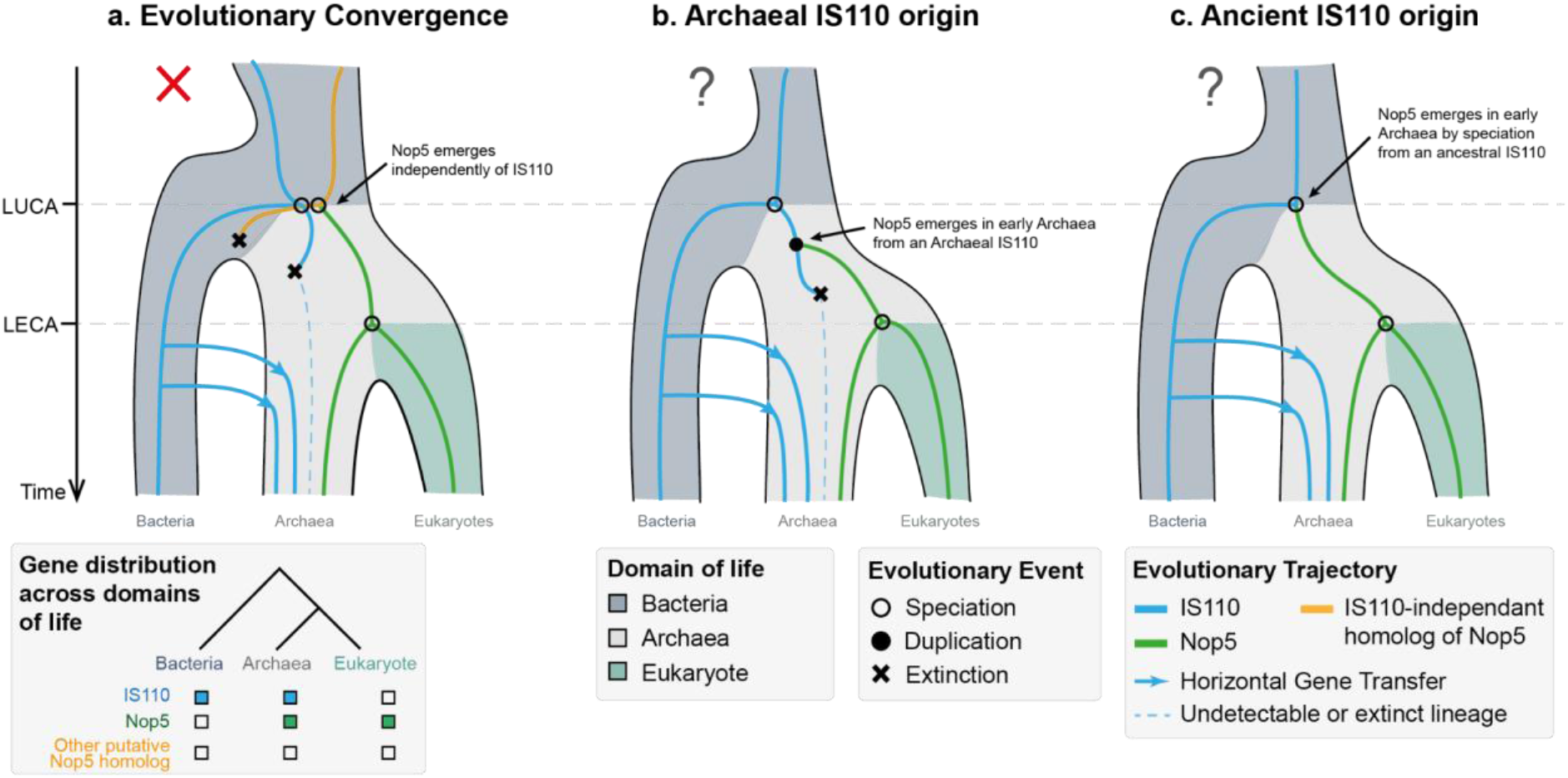
Three potential evolutionary scenarios to explain the relationship between IS110 Rec and Nop5 proteins. Each diagram describes a potential evolutionary trajectory of IS110 Rec (blue lines) and Nop5 (green lines) proteins among bacteria, archaea and eukaryotes (dark blue, gray and green colors) over time. The three scenarios explore distinct events, namely speciation (empty circle), duplication (filled circle), and extinction (cross). Note that all scenarios depict multiple lineages of archaeal IS110 Rec emerging via independent horizontal gene transfer (HGT) events, as supported by our phylogenetic analyses (**Supplementary Fig. 6**). **a,** The “Evolutionary Convergence” scenario posits that the observed structural similarity between IS110 Rec and Nop5 are not the consequence of shared ancestry but instead due to evolutionary convergence, leading to the association of similar domains. This scenario implies that the Nop5 lineage (green lines) emerged from an ancestor that was wholly independent of IS110 Rec (blue line), and emerged relatively early, given the widespread presence of Nop5 among archaea. We reject the “Convergence” scenario based on the following observations: (i) We were unable to detect IS110 Rec-independent homologs of Nop5 in our database, suggesting that if this ancestor once existed, it must have gone extinct; (ii) The conservation of key amino acid residues within Nop domain of IS110 Rec and Nop5 proteins renders a convergence scenario unlikely (**Supplementary Fig. 4a** and **b**); and (iii) IS110 Rec and Nop5 exist as sister clades within the broader family of RuvC-like proteins (Fig. 2a), indicating that the closest relative of Nop5 is likely to be IS110 Rec. **b,** The “Archaeal IS110 origin” scenario posits that the Nop5 lineage originated from an archaeal IS110 Rec through a duplication event (black circle). Under this scenario, one would expect to observe the whole clade of archaeal Nop5 and eukaryotic Nop56/Nop58 homologs to branch within a clade of archaeal IS110 Rec, however full-length and domain-specific phylogenetic trees do not exhibit such topologies. However, we cannot reject the possibility that the clade of archaeal IS110 Rec that gave rise to the Nop5 lineage have gone extinct in the past nor can we exclude that the clade of archaeal IS110 Rec homologs ancestral to Nop5 are not present in the archaeal genomes included in our database. Thus, we cannot completely reject this evolutionary scenario. **c,** The “Ancient IS110 origin” scenario posits that Nop5-family proteins and IS110 Rec are homologs and share a more ancient common ancestor. In the last common ancestor of bacteria and archaea, an ancestral protein would have given rise to both the IS110 Rec lineages in bacteria, and to Nop5 lineages in archaea (empty circle). This scenario is supported by the observed taxonomic distribution of proteins, the detection of IS110 Rec and Nop5 homologs across domains of life, and the topology of our phylogenetic trees showing that IS110 Rec and Nop5 proteins are sister clades within the family of RuvC-like proteins. We thus favor this evolutionary scenario for the emergence of the Nop5 lineage from its last common ancestor with the IS110 Rec lineage. The presence of ancestral features in IS110 Rec such as the DEDD catalytic motif, also found the ancient RuvC nuclease family of proteins, makes it more likely that the last common ancestor of Nop5 and IS110 Rec resembled IS110 Rec found in extant organisms.

**Supplementary Figure 6.**
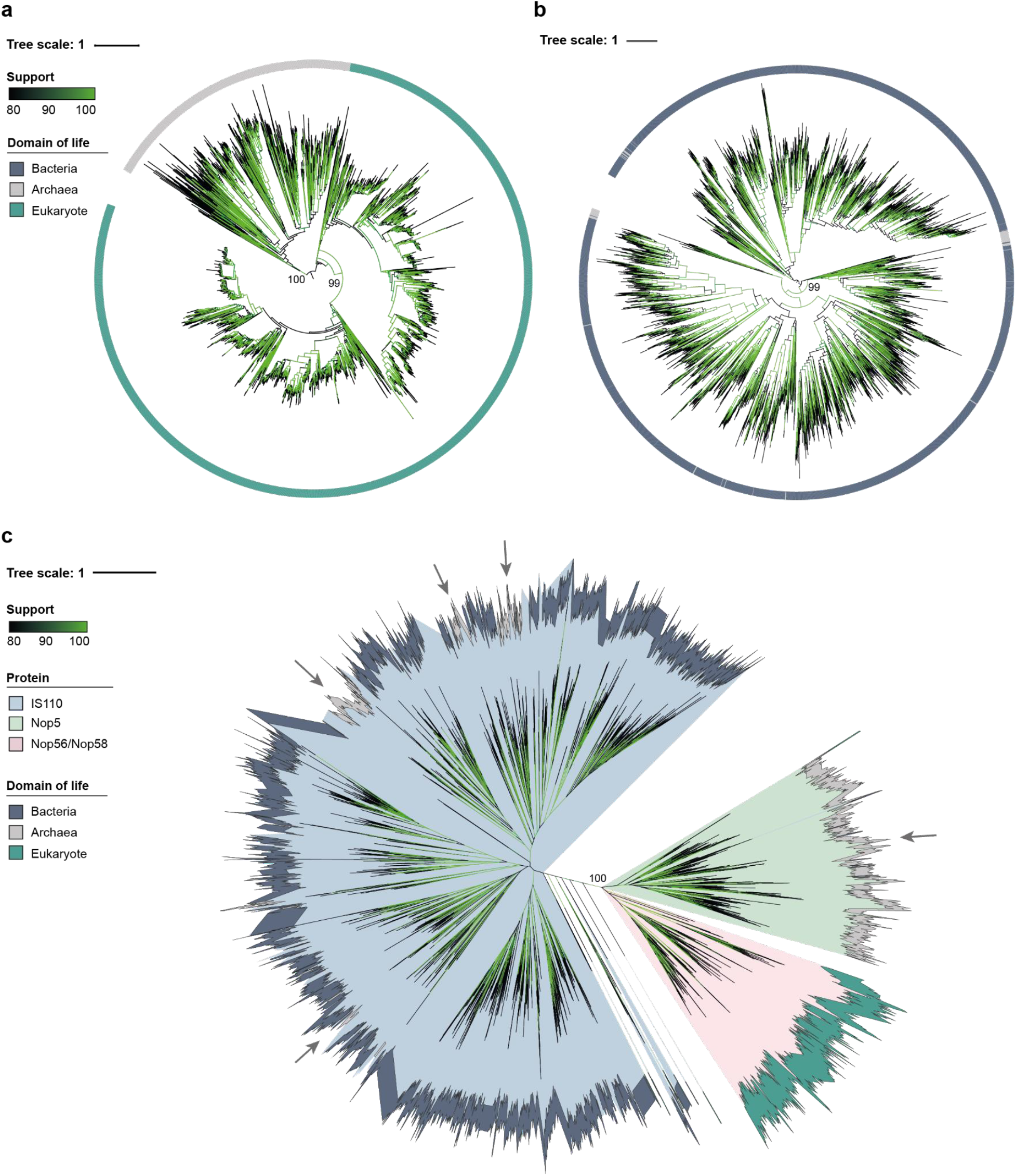
Phylogenetic tree based on full-length IS110 Rec, Nop5 and Nop56/Nop58 proteins. **a,** Phylogenetic tree of Nop5 and Nop56/Nop58 homologs built on full-length protein sequences. Archaeal Nop5 homologs were chosen as an outgroup to root the tree, and the outer ring indicates the domain of life that each protein belongs to. UltraFast Bootstrap support values are represented in shades of green on the tree branches, and explicit values are indicated for key branches. **b,** Phylogenetic tree of IS110 Rec homologs, built on full-length protein sequences, colored as in **a**. The tree is midpoint rooted. **c,** Unrooted phylogenetic tree of IS110 Rec and Nop5 protein homologs aligned on their full-length amino acid sequence. Inner ring colors indicate the protein identity, and outer ring colors indicate which domain of life each protein belongs to; UltraFast Bootstrap support values are indicated in shades of green on the tree branches. Gray arrows indicate major archaeal clades in the tree. IS110 Rec homologs found in archaeal organisms are distributed across several distinct clades and are mixed with bacterial homologs. This observation strongly supports the idea that archaeal IS110 transposable elements were acquired in archaea through multiple independent horizontal gene transfer (HGT) events.

**Supplementary Figure 7.**
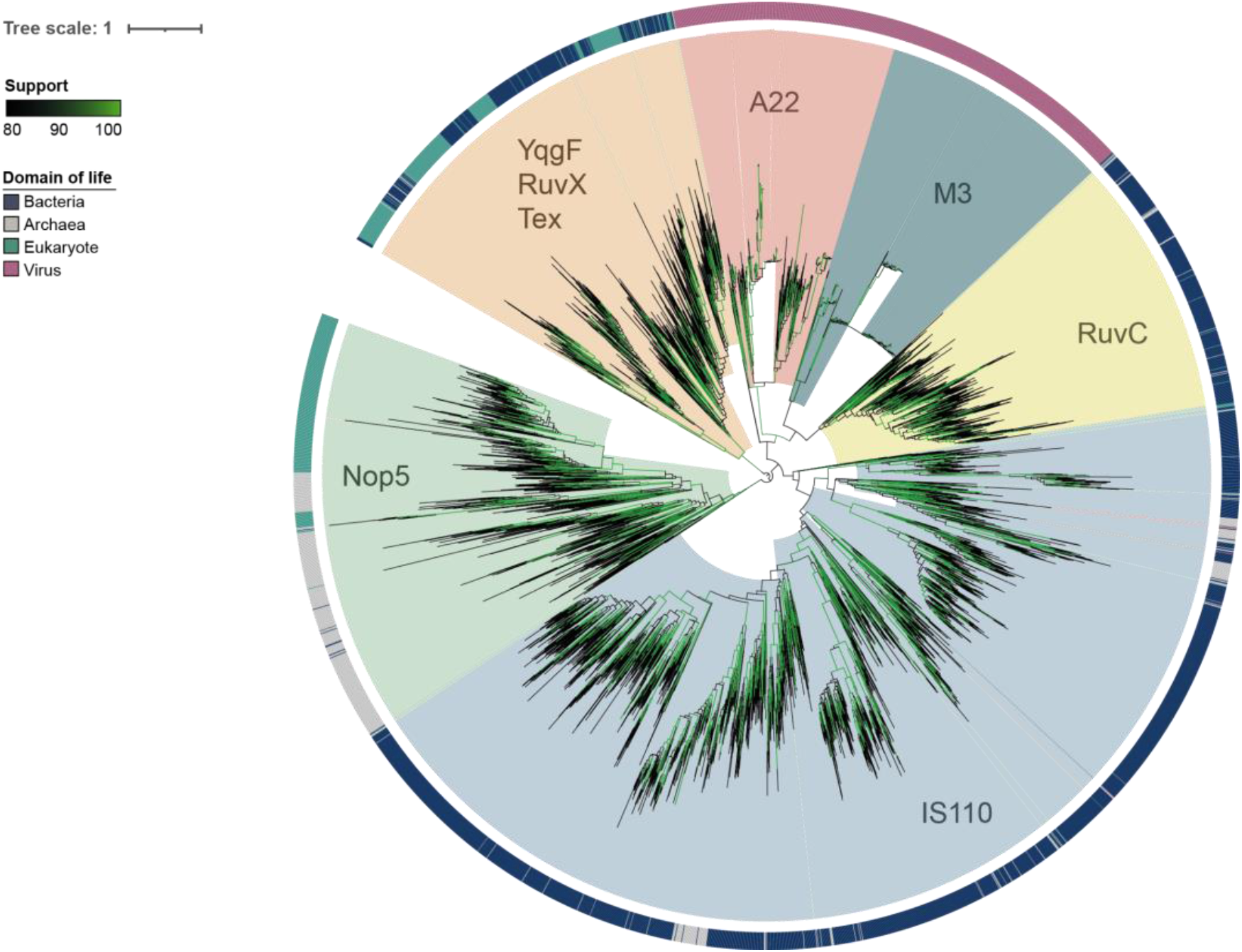
Phylogenetic tree built on the N-terminal domain of IS110 Rec, Nop5, and Nop56/Nop58. Candidate homologs were identified by querying a multiple sequence alignment of the N-terminal domain of IS110 Rec, Nop5, and Nop56/Nop58, alongside RuvC Holliday Junction Resolvase proteins, in HHpred against PfamA-v36 (see **Methods**). The YqgF/RuvX/Tex clade of proteins was chosen as an outgroup to root the tree (orange). UltraFast Bootstrap support values for each branch are indicated in green and are explicitly specified for key branches.

## REFERENCES

1 Martin, E. C. et al. Insights into RAG Evolution from the Identification of “Missing Link” Family A RAGL Transposons. Mol Biol Evol 40 (2023). 10.1093/molbev/msad232

2 Michelini, F. et al. From “Cellular” RNA to “Smart” RNA: Multiple Roles of RNA in Genome Stability and Beyond. Chem Rev 118, 4365–4403 (2018). 10.1021/acs.chemrev.7b00487

3 Dlakić, M. & Mushegian, A. Prp8, the pivotal protein of the spliceosomal catalytic center, evolved from a retroelement-encoded reverse transcriptase. Rna 17, 799–808 (2011). 10.1261/rna.2396011

4 Podlevsky, J. D. & Chen, J. J. Evolutionary perspectives of telomerase RNA structure and function. RNA Biol 13, 720–732 (2016). 10.1080/15476286.2016.1205768

5 Altae-Tran, H. et al. The widespread IS200/IS605 transposon family encodes diverse programmable RNA-guided endonucleases. Science (New York, N.Y.) 374, 57–65 (2021). 10.1126/science.abj6856

6 Karvelis, T. et al. Transposon-associated TnpB is a programmable RNA-guided DNA endonuclease. Nature 599, 692–696 (2021). 10.1038/s41586-021-04058-1

7 Meers, C. et al. Transposon-encoded nucleases use guide RNAs to promote their selfish spread. Nature 622, 863–871 (2023). 10.1038/s41586-023-06597-1

8 Yu, G., Zhao, Y. & Li, H. The multistructural forms of box C/D ribonucleoprotein particles. Rna 24, 1625–1633 (2018). 10.1261/rna.068312.118

9 Kiss, T. Small nucleolar RNAs: an abundant group of noncoding RNAs with diverse cellular functions. Cell 109, 145–148 (2002). 10.1016/s0092-8674(02)00718-3

10 Monaco, P. L., Marcel, V., Diaz, J. J. & Catez, F. 2’-O-Methylation of Ribosomal RNA: Towards an Epitranscriptomic Control of Translation? Biomolecules 8 (2018). 10.3390/biom8040106

11 Wilkinson, M. E., Charenton, C. & Nagai, K. RNA Splicing by the Spliceosome. Annu Rev Biochem 89, 359–388 (2020). 10.1146/annurev-biochem-091719-064225

12 Liu, S. et al. Binding of the human Prp31 Nop domain to a composite RNA-protein platform in U4 snRNP. Science (New York, N.Y.) 316, 115–120 (2007). 10.1126/science.1137924

13 Liu, S., Ghalei, H., Lührmann, R. & Wahl, M. C. Structural basis for the dual U4 and U4atac snRNA-binding specificity of spliceosomal protein hPrp31. Rna 17, 1655–1663 (2011). 10.1261/rna.2690611

14 Koonin, E. V., Bork, P. & Sander, C. A novel RNA-binding motif in omnipotent suppressors of translation termination, ribosomal proteins and a ribosome modification enzyme? Nucleic Acids Res 22, 2166–2167 (1994). 10.1093/nar/22.11.2166

15 Choi, S., Ohta, S. & Ohtsubo, E. A novel IS element, IS621, of the IS110/IS492 family transposes to a specific site in repetitive extragenic palindromic sequences in Escherichia coli. J Bacteriol 185, 4891–4900 (2003). 10.1128/jb.185.16.4891-4900.2003

16 Söding, J., Biegert, A. & Lupas, A. N. The HHpred interactive server for protein homology detection and structure prediction. Nucleic Acids Res 33, W244–248 (2005). 10.1093/nar/gki408

17 Wan, R. et al. The 3.8 Å structure of the U4/U6.U5 tri-snRNP: Insights into spliceosome assembly and catalysis. Science (New York, N.Y.) 351, 466–475 (2016). 10.1126/science.aad6466

18 Lapinaite, A. et al. The structure of the box C/D enzyme reveals regulation of RNA methylation. Nature 502, 519–523 (2013). 10.1038/nature12581

19 Durrant, M. G. et al. Bridge RNAs direct modular and programmable recombination of target and donor DNA. bioRxiv (2024). 10.1101/2024.01.24.577089

20 Rezwan Siddiquee, C. H. P., Ruth M. Hall, Sandro F. Ataide. A programmable seeker RNA guides target selection by IS1111 and IS110 type insertion sequences. bioRxiv (2024). doi: 10.1101/2024.04.26.591405

21 Jumper, J. et al. Highly accurate protein structure prediction with AlphaFold. Nature 596, 583–589 (2021). 10.1038/s41586-021-03819-2

22 Wang, J., Yang, Z. & Ye, K. Methylation guide RNAs without box C/D motifs. Rna 28, 1597–1605 (2022). 10.1261/rna.079379.122

23 Ye, K. et al. Structural organization of box C/D RNA-guided RNA methyltransferase. Proc Natl Acad Sci U S A 106, 13808–13813 (2009). 10.1073/pnas.0905128106

24 Ghalei, H., Hsiao, H. H., Urlaub, H., Wahl, M. C. & Watkins, N. J. A novel Nop5-sRNA interaction that is required for efficient archaeal box C/D sRNP formation. Rna 16, 2341–2348 (2010). 10.1261/rna.2380410

25 Stark, W. M. The Serine Recombinases. Microbiol Spectr 2 (2014). 10.1128/microbiolspec.MDNA3-0046-2014

26 van Kempen, M. et al. Fast and accurate protein structure search with Foldseek. Nature biotechnology 42, 243–246 (2024). 10.1038/s41587-023-01773-0

27 Zimmermann, L. et al. A Completely Reimplemented MPI Bioinformatics Toolkit with a New HHpred Server at its Core. J Mol Biol 430, 2237–2243 (2018). 10.1016/j.jmb.2017.12.007

28 Majorek, K. A. et al. The RNase H-like superfamily: new members, comparative structural analysis and evolutionary classification. Nucleic Acids Res 42, 4160–4179 (2014). 10.1093/nar/gkt1414

29 Aravind, L., Makarova, K. S. & Koonin, E. V. SURVEY AND SUMMARY: holliday junction resolvases and related nucleases: identification of new families, phyletic distribution and evolutionary trajectories. Nucleic Acids Res 28, 3417–3432 (2000). 10.1093/nar/28.18.3417

30 Feng, J. M., Tian, H. F. & Wen, J. F. Origin and evolution of the eukaryotic SSU processome revealed by a comprehensive genomic analysis and implications for the origin of the nucleolus. Genome biology and evolution 5, 2255–2267 (2013). 10.1093/gbe/evt173

31 Kapitonov, V. V., Makarova, K. S. & Koonin, E. V. ISC, a Novel Group of Bacterial and Archaeal DNA Transposons That Encode Cas9 Homologs. J Bacteriol 198, 797–807 (2015). 10.1128/jb.00783-15

32 Chylinski, K., Makarova, K. S., Charpentier, E. & Koonin, E. V. Classification and evolution of type II CRISPR-Cas systems. Nucleic Acids Res 42, 6091–6105 (2014). 10.1093/nar/gku241

33 Sayers, E. W. et al. GenBank. Nucleic Acids Res 47, D94–d99 (2019). 10.1093/nar/gky989

34 Parks, D. H. et al. GTDB: an ongoing census of bacterial and archaeal diversity through a phylogenetically consistent, rank normalized and complete genome-based taxonomy. Nucleic Acids Res 50, D785–d794 (2022). 10.1093/nar/gkab776

35 Mistry, J. et al. Pfam: The protein families database in 2021. Nucleic Acids Res 49, D412–d419 (2021). 10.1093/nar/gkaa913

36 Varadi, M. et al. AlphaFold Protein Structure Database: massively expanding the structural coverage of protein-sequence space with high-accuracy models. Nucleic Acids Res 50, D439–d444 (2022). 10.1093/nar/gkab1061

37 Berman, H. M. et al. The Protein Data Bank. Nucleic Acids Res 28, 235–242 (2000). 10.1093/nar/28.1.235

38 Mirdita, M. et al. ColabFold: making protein folding accessible to all. Nat Methods 19, 679–682 (2022). 10.1038/s41592-022-01488-1

39 Evans, R. O. N. M.; Pritzel, A.; Antropova, N.; Senior, A.; Green, T.; Žídek, A.; Bates, R.; Blackwell, S.; Yim, J.; Ronneberger, O.; Bodenstein, S.; Zielinski, M.; Bridgland, A.; Potapenko, A.; Cowie, A.; Tunyasuvunakool, K.; Jain, R.; Clancy, E.; Kohli, P.; Jumper, J.; Hassabis, D. Protein complex prediction with AlphaFold-Multimer. bioRxiv (2022). 10.1101/2021.10.04.463034

40 Meng, E. C. et al. UCSF ChimeraX: Tools for structure building and analysis. Protein Sci 32, e4792 (2023). 10.1002/pro.4792

41 Altschul, S. F., Gish, W., Miller, W., Myers, E. W. & Lipman, D. J. Basic local alignment search tool. J Mol Biol 215, 403–410 (1990). 10.1016/s0022-2836(05)80360-2

42 Katoh, K., Misawa, K., Kuma, K. & Miyata, T. MAFFT: a novel method for rapid multiple sequence alignment based on fast Fourier transform. Nucleic Acids Res 30, 3059–3066 (2002). 10.1093/nar/gkf436

43 Eddy, S. R. Accelerated Profile HMM Searches. PLoS Comput Biol 7, e1002195 (2011). 10.1371/journal.pcbi.1002195

44 Hauser, M., Steinegger, M. & Söding, J. MMseqs software suite for fast and deep clustering and searching of large protein sequence sets. Bioinformatics 32, 1323–1330 (2016). 10.1093/bioinformatics/btw006

45 Protsyuk, I. V., Grekhov, G. A., Tiunov, A. V. & Fursov, M. Y. Shared bioinformatics databases within the Unipro UGENE platform. J Integr Bioinform 12, 257 (2015). 10.2390/biecoll-jib-2015-257

46 Sayers, E. W. et al. Database resources of the National Center for Biotechnology Information. Nucleic Acids Res 52, D33–d43 (2024). 10.1093/nar/gkad1044

47 Biegert, A., Mayer, C., Remmert, M., Söding, J. & Lupas, A. N. The MPI Bioinformatics Toolkit for protein sequence analysis. Nucleic Acids Res 34, W335–339 (2006). 10.1093/nar/gkl217

48 Steenwyk, J. L., Buida, T. J., 3rd, Li, Y., Shen, X. X. & Rokas, A. ClipKIT: A multiple sequence alignment trimming software for accurate phylogenomic inference. PLoS Biol 18, e3001007 (2020). 10.1371/journal.pbio.3001007

49 Nguyen, L. T., Schmidt, H. A., von Haeseler, A. & Minh, B. Q. IQ-TREE: a fast and effective stochastic algorithm for estimating maximum-likelihood phylogenies. Mol Biol Evol 32, 268–274 (2015). 10.1093/molbev/msu300

50 Hoang, D. T., Chernomor, O., von Haeseler, A., Minh, B. Q. & Vinh, L. S. UFBoot2: Improving the Ultrafast Bootstrap Approximation. Mol Biol Evol 35, 518–522 (2018). 10.1093/molbev/msx281

51 Kalyaanamoorthy, S., Minh, B. Q., Wong, T. K. F., von Haeseler, A. & Jermiin, L. S. ModelFinder: fast model selection for accurate phylogenetic estimates. Nat Methods 14, 587–589 (2017). 10.1038/nmeth.4285

